# An evolutionary algorithm for inverse RNA folding inspired by Lévy flights

**DOI:** 10.1101/2022.01.17.476593

**Authors:** Nono S. C. Merleau, Matteo Smerlak

**Affiliations:** Max Planck Institute for Mathematics in the Sciences, Leipzig, Germany

## Abstract

A Lévy flight is a random walk with step sizes that follow a heavy-tailed probability distribution. This type of random walk, with many small steps and a few large ones, has inspired many applications in genetic programming and evolutionary algorithms in recent years, but is yet to be applied to RNA design. Here we study the inverse folding problem for RNA, viz. the discovery of sequences that fold into given target secondary structures. We implement a Lévy mutation scheme in an updated version of `aRNAque`, an evolutionary inverse folding algorithm, and apply it to the design of RNAs with and without pseudoknots. We find that the Lévy mutation scheme increases the diversity of designed RNA sequences and reduces the average number of evaluations of the evolutionary algorithm. The results show improved performance on both `Pseudobase++` and the `Eterna100` datasets, outperforming existing inverse folding tools. We propose that a Lévy flight offers a better standard mutation scheme for optimizing RNA design.

## Introduction

The function of non-coding RNA, which includes gene expression regulation (miRNAs, piRNAs, lncRNAs), RNA maturation (snRNAs, snoRNAs) and protein synthesis (rRNAs, tRNAs), strongly depends on the hierarchical folding of RNA molecules. Given their sequence of bases (primary structure), RNAs fold into secondary structures, such as stem loops and pseudoknots, before folding into higher level (tertiary and quaternary) structures. The secondary structure can be considered as a list of base pairs, including canonical, Watson-Crick pairs (1, 2); non-canonical interactions, that occur with reduced frequency; and crossing or pseudo-knot interactions (when two canonical or non-canonical interactions cross each other) (3).

Here, we consider the RNA secondary structure inverse folding problem. The goal is to find RNA sequences that fold into a given target secondary structure, with or without pseudoknots. Considering pseudoknots in designing functional RNAs is vital given their role in realising biological functions. In modern bio-engineering, one must solve the RNA inverse folding problem to be able to design RNA molecules performing specific functions (4–6).

A key prerequisite to addressing the RNA inverse problem is a reliable solution to the folding problem. Computationally folding an RNA molecule consists of searching in the space of all possible secondary structures for one that minimises the free energy. Designing sequences for a pseudo-knotted target structure is computationally more expensive than a pseudoknot-free target because of the complexity of the folding algorithms required. Specifically, the time complexity of the pseudoknot-free secondary structure prediction is *O*(*n*^3^) when using dynamic programming approaches such as `RNAfold` (7), or less with heuristic folding methods (e.g. *O*(*n*) for `LinearFold` and *O*(*n*^2^ log *n*) for `RAFFT` (8)). By contrast, when considering a special class of pseudoknots, the time complexity of folding goes up to *O*(*n*^6^) for an exact thermodynamic prediction using a dynamic programming approach such as (9). In this work, we consider only two heuristics tools (IPknot (10) and `HotKnots` (11)) chosen for their lower time complexity *O*(*n*^4^).

Many of the studies addressing the inverse folding of RNA considered only pseudoknot-free secondary structures (12–19). There are, however, three exceptions, the most recent of which is `antaRNA` (20) utilising the “ant-colony” optimisation technique. The technique begins with an initial sequence generated via a weighted random search; next the solutions are evaluated, and the sequence fitness values are used to refine the weights and improve the sequences over generations. Another approach (`Modena`) implements a multi-objective evolutionary algorithm measuring both the stability of the designed sequences and the similarities of folded sequences to the target structure. Although the first version of `Modena` was implemented for pseudoknot-free structure (15), it has since been extended to support pseudoknotted RNAs and a new crossover operator (21). `Inv` (22) is the first inverse folding tool handling pseudoknotted RNA target structures, but is restricted to a specific type of pseudoknot pattern called 3-crossing nonplanar pseudoknots. In a prior publication (23), we presented `aRNAque`, a simple evolutionary inverse folding algorithm guided by local (or one-point mutations). Although a local search can efficiently discover optima in a simple landscape, more complex landscapes pose challenges to the design of evolutionary algorithms that rely solely on local search. This is especially true on a neutral landscape where local search may be inefficient or risk getting stuck on a plateau (or local optimum). To avoid this pitfall, we propose here an extension of `aRNAque` which implements a new mutation scheme inspired by Lévy flights (called Lévy mutation) and supports pseudoknotted RNA target structures.

Lévy flights are random walks with a heavy-tailed step size distribution. The concept originates in the work of Mandelbrot on the fluctuation of commodities prices in the 1960s (24), but has since found many more physical applications (25). Lévy flights also play a key role in the context of animal foraging, perhaps because they provide an optimal balance between exploration and exploitation (26, 27). For a recent review of applications of Lévy flights in biology from the molecular to the ecological scale (28).

Similar to a Lévy flight, a Lévy mutation scheme allows simultaneous search at all scales over the landscape. New mutations most often produce nearby sequences (one-point mutations), but occasionally generate mutant sequences which are far away in genotype space (macro-mutations). In this work, the distribution of the number of point mutations at every step is taken to follow a Zipf distribution (29).

The optimization approach implemented in `aRNAque` is an evolutionary algorithm, which consists of a population of RNA sequences that all perform separate random walks (are mutated) in the space of possible sequences, and whose stepsizes (number of point mutations) follow a Zipf distribution. After each step, the probability of surviving is proportional to the fitness of each sequence, which is evaluated by its ability to approximate a given target structure. We provide a brief overview of that approach in the following subsection.

Earlier works have applied similar ideas in genetic programming (30), and in differential evolutionary algorithms (31). This has motivated us to investigate the possible benefit of a Lévy flight in the design of RNA sequences. Using a Lévy mutation scheme, we aim to speed up our prior evolutionary algorithm and increase the diversity of the designed RNA sequences.

We compared the performance of our newly modified version of `aRNAque` to existing tools through a benchmark on two well-known RNA datasets: `PseudoBase+++` (32) for the pseudoknotted targets and `Eterna100` (33) for the pseudoknot-free targets. On the `PseudoBase+++` dataset, the difference between the local mutation and the Lévy mutation with respect to the number of generations (or evaluations) was significant (with a *p*-value ≈0.00004). Using the two pseudoknot folding tools `HotKnots` and `IPknot`, our designed sequences were of better quality than the ones produced by `antaRNA` regarding the average base pair distance to the desired targets. We performed a second benchmark on the `Eterna100` dataset. Considering the `Eterna100-V1` dataset, the Lévy mutation scheme solved 89 targets out of 100 whereas the local mutation scheme solved 91*/*100. To compare `aRNAque` to the existing pseudoknot-free inverse folding tools, we combined the two benchmark results obtained using both mutation schemes, and we counted the total number of distinct targets solved. `aRNAque` designs successfully 92*/*100 of the `Eterna100-V1` dataset and 94*/*100 of the `Eterna100-V2` dataset.

### Evolutionary algorithm (EA)

Below, we provide a brief overview of our evolutionary search algorithm and our mutation scheme.

#### Overview

In general, an evolutionary search algorithm on any fitness landscape consists of three main parts, which in the context of RNA inverse folding are as follows:

- Initialization: generating a random initial population of RNA sequences compatible with the given target secondary structure.
- Evaluation and selection: evaluating a population of RNA sequences consists of two steps: 1) fold each sequence into a secondary structure and assign it a weight based on its similarity to the target structure. 2) select a weighted random sample with replacement from the current population to generate a new population. A detailed description of the objective function used in `aRNAque` is provided in (23).
- Mutation (or move) operation: define a set of rules or steps used to produce new sequences from the selected or initial ones. This component is elaborated further in the next subsection.

#### Mutation mode

For a given target RNA secondary structure *σ*\* of length *L*, the space of potential solutions to the inverse folding problem is *S* = {*A, C, G, U*} ^*L*^. An evolutionary algorithm explores the space *S* through its move (or mutation) operator.

Given a sequence *ϕ* ∈ *S*, a sequence *ϕ*^′^ ∈ *S* is said to be an *n*-point mutation of *ϕ* if it differs from *ϕ* at *n* nucleotides; i.e. *h*(*ϕ, ϕ*^′^) = *n* where *h*(.,.) is the hamming distance on *S*.

A mutation mode is a random variable *U* taking values in 1, …, *L*. *P* (*U* = *n*) is defined as the probability that, exactly *n* nucleotides, selected uniformly at random undergo point mutation during a mutation event. *U* can generally be any probability distribution. We examined the binomial and Zipf distributions for local and Levy search, respectively:

- Binomial mutation: *U* has a binomial distribution:

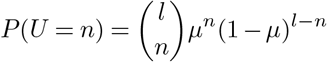

for some 0 ≤ *μ* ≤ 1, such that *u* = *μ* · *l*. We can think of this mutation mode arising from each nucleotide of an RNA sequence independently undergoing a point mutation with probability *μ*, i.e. *μ* is the per-nucleotide or point mutation rate.
- Lévy mutation: *U* has a Zipf distribution given by:

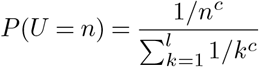

where *c* > 0 is the value of the exponent characterizing the distribution. Larger values of *c* are associated a greater proportion of local search, while smaller values of *c* imply a greater proportion of long-range search.

Figure 1 shows the distribution of the number of point mutations on a sequence of length 88 nucleotides for both mutation schemes. Both distributions have the same mean, but differ markedly in their tails.

**Fig. 1.**
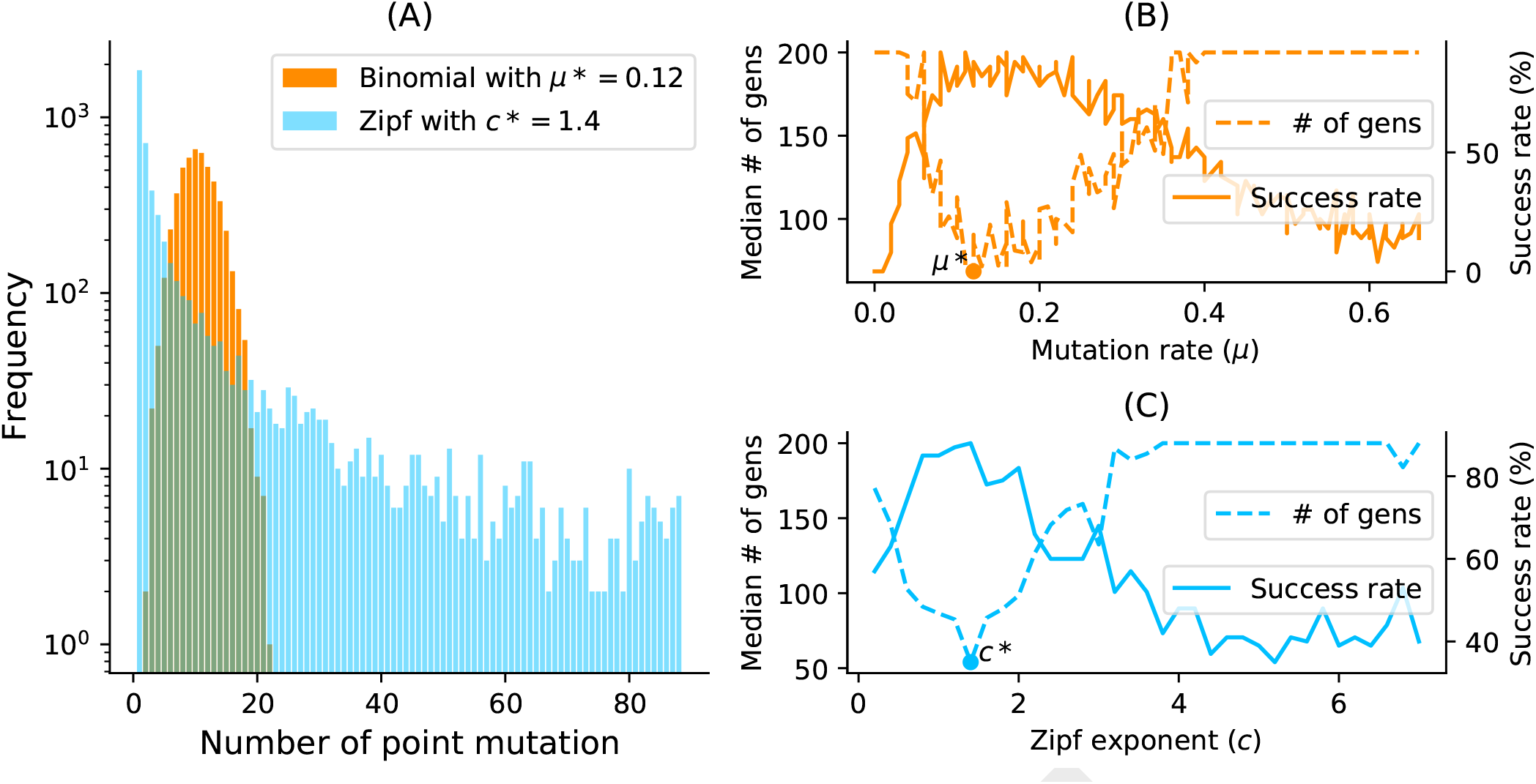
(A) Samplings Binomial and Zipf distributions for the best binomial mutation rate *μ*^*\**^ (respectively *c*^*\**^ for the best Zipf exponent parameter). Both distributions have a mean of 8.7 point mutations for a sequence of length 88 nucleotides. (B) Tuning of binomial mutation rate parameter. For each *μ* ∈ [0, 1] with a step size of 0.005 and the pseudoknotted target `PKB00342` of length 88, 50 sequences were designed using `aRNAque`. (B) shows the median generations and the success percentage *vs*. the mutation rate (*μ*). The best mutation rate is *μ*^*\**^ = 0.085 (with a median number of generation 93.5 and a success rate of 92%). (C) Tuning of Levy exponent. Similar to (B), for each *c* ∈ [0, 7] with a step size of 0.1 and for the same pseudoknotted target structure, 100 sequences were designed using `aRNAque`. It shows the median generations and the percentage of success *vs*. the exponent parameter (*c*). The Zipf exponent distribution that produced the highest success rate and the minimum number of generations is *c*^*\**^ = 1.4..

##### Algorithm 1: Mutation algorithm

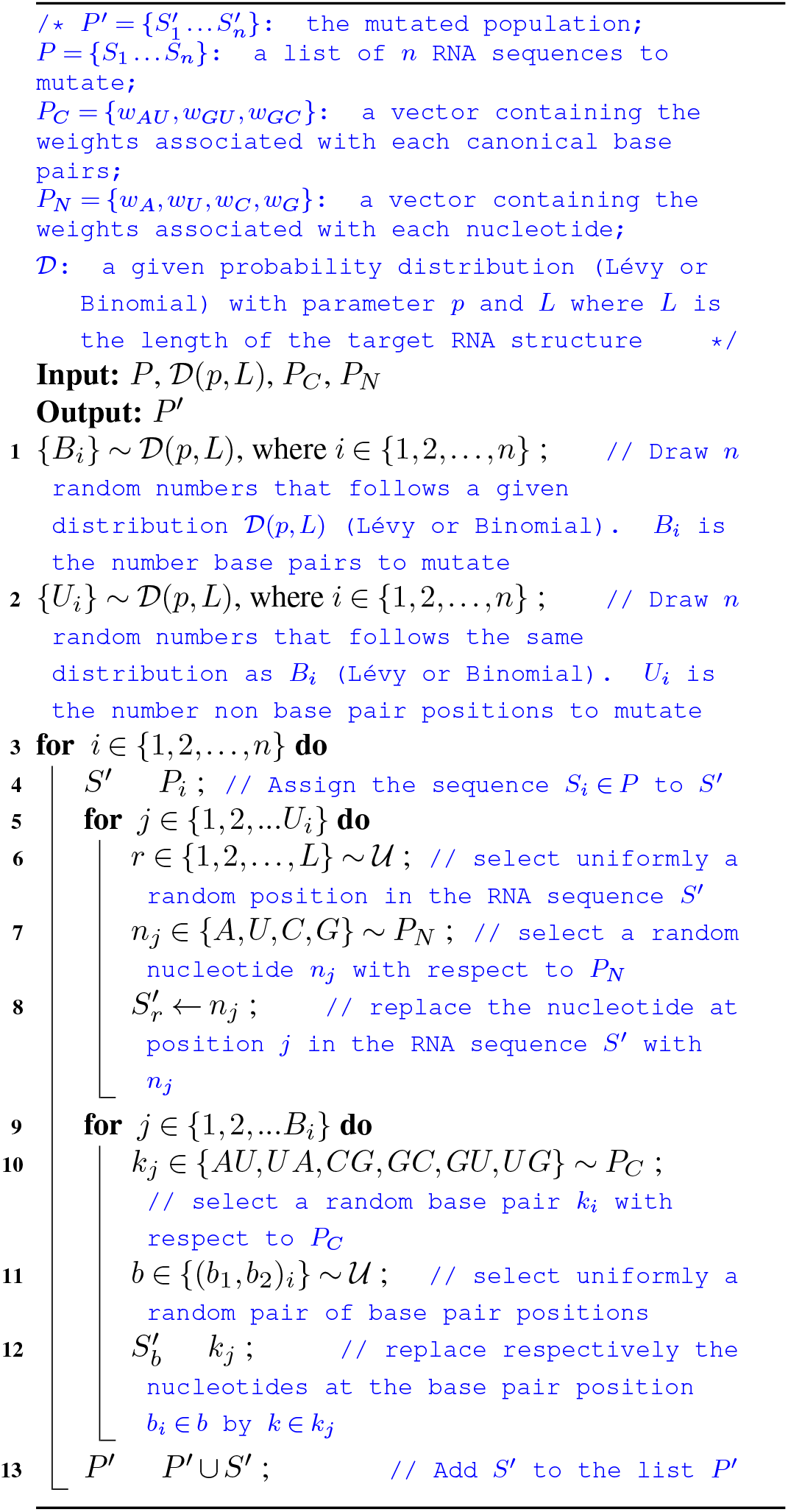

Throughout this paper, local mutation will refer to binomial distributed mutation with parameter *μ* ≈ 1*/L* or to one-point mutation.

#### New feature

We provide an updated version of `aRNAque` supporting pseudoknotted RNA target structures. In addition to the support for pseudoknots, we provide an updated mutation mode based on a Zipf distribution. We present the mutation algorithm in Algorithm 1.

#### Parameter analysis and benchmark

Here we analyse mutation parameters and compare local and Lévy mutation modes.

#### Benchmark data

To compare our new version of `aRNAque` with existing tools in the literature, we used the `PseudoBase++` benchmark datasets for pseudoknotted target structures and the `Eterna100` dataset for pseudoknot-free target structures.

The `PseudoBase++` is a set of 265 pseudoknotted RNA structures used to benchmark `Modena`. It was initially 304 RNA secondary structures, but we excluded 37 because they had non-canonical base pairs. We then grouped the structures into four pseudoknot motifs (Figure 2): 209 hairpin pseudoknots (H), 29 bulge pseudoknots (B), 8 complex hairpin pseudoknots (cH) and 4 kissing hairpin pseudoknots (K).

**Fig. 2.**
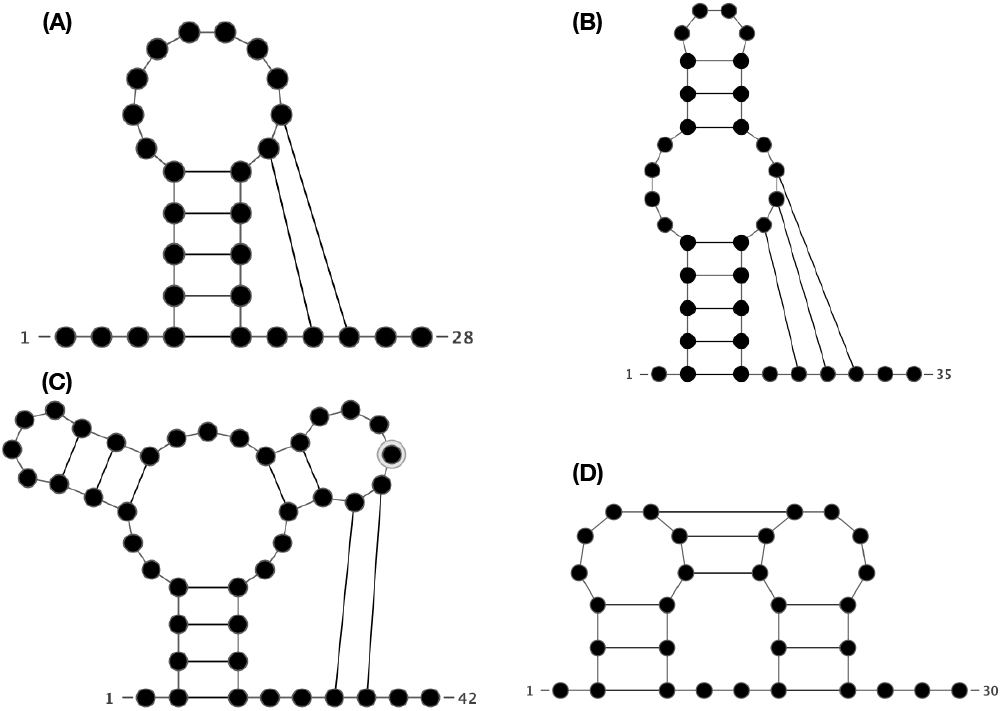
Types of pseudoknots accommodated by aRNAque. (A) Hairpin (H-type) pseudoknot. (B) Bulge (B-type) pseudoknot. (C) Complex hairpin (cH-type) pseudoknot. (D) Kissing hairpin (K-type) pseudoknot.

The `Eterna100` dataset (34) is available in two versions and both contain a set of 100 target structures extracted from the `EteRNA` puzzle game and classified by their degree of difficulty. The `Eterna100-V1` was initially designed using `ViennaRNA` 1.8.5, which relies on Turner1999 energy parameters (35). Out of the 100 target secondary structures, 19 turned out to be unsolvable using the recent version of `ViennaRNA` (Version 2.14). Subsequently, an `Eterna100-V2` (34) was released in which the 19 targets were slightly modified to be solvable using `ViennaRNA 2.14`.

#### Methodology

The best mutation parameters obtained for both binomial and Lévy mutation modes are used to benchmark and compare the results on the entire datasets of RNA structures (265 from `PseudoBase++` and 100 from `EteRNA100`). First, for each of the 365 target structures *σ*^*^ in the datasets, 20 sequences were designed. To measure the performance of each tool, each designed sequence *s* is folded into a secondary structure *σ* and the similarities between *σ* and *σ*^*^ are computed using the base pair distance. Second, for each of the `Eterna100` target structures and a maximum of 5000 generations (i.e. 50, 000 evaluations), 5 to 20 runs were launched independently, which results in at least 5 designed sequences per target. We define success rate simply as the number of successfully designed targets. A target is considered successfully designed when at least one of the designed sequence folds into the target structure (i.e. the Hamming distance between the target structure and the MFE structure is 0).

##### Folding tools

Two tools for pseudoknotted RNA folding are considered in this work: `HotKnots` and `IPknot`. For pseudoknot-free RNA folding, we used `RNAfold`. For the mutation parameter analysis presented here, we used `IPknot`, and both `HotKnots` and `IPknot` for pseudoknotted targets. Furthermore, we considered `pkiss`, a well know tool for K-type pseudoknot prediction, but since the `PseudoBase++` dataset contains just 5 K-type pseudoknotted structures and `pKiss` has higher time complexity (*O*(*n*^6^)), we did not find it efficient for the benchmark we performed.

##### Mutation parameters tuning

One of the main challenges for an evolutionary algorithm is to find optimum parameters such as mutation rate, population size and selection function. We used 81 pseudoknotted targets with lengths from 25 to 181 nucleotides for the mutation parameter analysis. We set the maximum number of generations to 200 and the population size to 100. The best mutation parameters (*c*^*^ for Levy and *μ*^*^ for Binomial) are those that have the lowest median number of generations.

- Binomial mutation: From Figure 1B, the critical range was identified to be from 0 to 0.2 and as *μ* becomes greater than 0.1, the success rate decreases and the average number of generations increases. For each of the 80 target structures with pseudoknots, 20 sequences were designed for *μ* ∈ [0, 0.2] with a step size of 1*/L*. Figure 3B shows the histogram of the best mutation rate found for each target structure. Two main regimes are apparent: one where the best mutation rate is very low mutation rate (≈1*/L*) and another where the high mutation rate is optimal.
- Lévy mutation: From Figure 1C, the critical range of *c* was identified to be [1, 2]. For *c* ∈ [1, 2] and a step size of 0.1, an optimum exponent parameter *c*^*^ was investigated for all the 80 target structures. Figure 3A shows the histogram of *c*^*^. Contrary to binomial mutation, the optimum exponent parameter does not vary too much (∀*σ, c*^*^ ≈ 1).

**Fig. 3.**
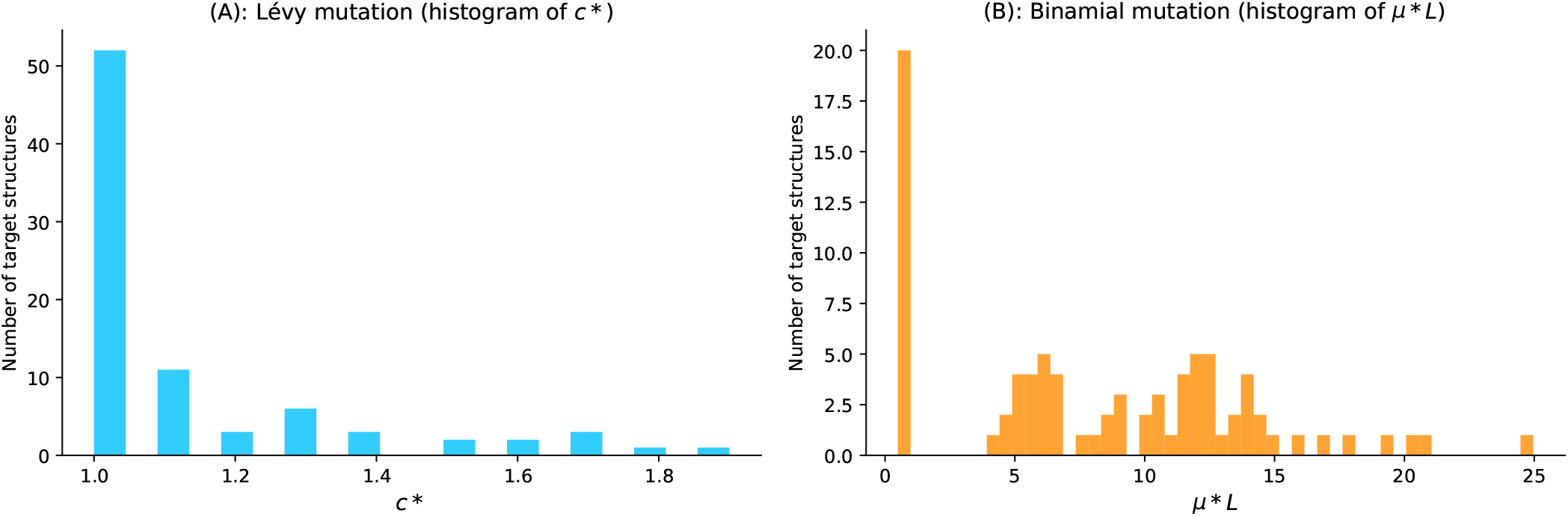
Parameter tuning for both binomial and Lévy mutation schemes.(A) Lévy mutation parameter tuning. Histogram of best exponent parameter (*c*^*\**^) for a set of 81 target structures with different pseudoknot patterns and various lengths. The most frequent best exponent value is 1. (B) Binomial parameter tuning. Histogram of best mutation rate (*μ*^*\**^) for the same set of 81 target structures with different pseudoknots and various lengths. The most frequent best parameter is the low mutation rate (≈ 1*/L*). For some structures, the best mutation rate is the high one for different lengths as well.

Figure 3A shows that when using a Lévy mutation, the optimal mutation rate is approximately the same for most target structures. In contrast, the optimum binomial mutation rate parameter *μ*^*^ mostly varies with different targets. In Figures 1B and 1C, although both mutation modes have approximately the same success rates (88% for the Lévy over 100 runs and ≈ 92% for the binomial over 50 runs), the Lévy flight mutation scheme is more robust to different targets. Moreover, the median number of generations for the Lévy mutation is lower (54 for the Lévy and 92 for the binomial mutation mode), thus enhancing efficiency.

## Results

We first compared the performance of `aRNAque` using Lévy mutations to the previous version with local mutations (binomial number of point-mutations with 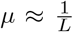). Secondly, we compared `aRNAque` to the existing pseudoknotted RNA inverse folding tool `antaRNA` using two folding tools: `HotKnots` and `IPknot`. We used the `PseudoBase++` dataset for both benchmarks.

### Performance on PseudoBase++: Levy mutation *vs*. local mutation

Figure 4 shows box plots for the base pair distance (Hamming distance) and the number of generations for increasing target lengths under our two mutation schemes: binomial at low mutation rate (or one point mutation) and the Lévy mutation. For each pseudoknotted RNA target structure in the `PseudoBase++` dataset, we designed 20 sequences. The results show that using the Lévy mutation instead of a local mutation scheme can significantly increase the performance of `aRNAque`. The gain was less significant in terms of designed sequences quality (base pair distance distributions, with a *t*-value ≈ − 1.04 and *p*-value ≈0.16) but more significant in terms of the average minimum number of generations needed for successful matches to target structures (with a *t*-value ≈ − 3.6 and *p*-value ≈ 0.0004). This result demonstrates a substantial gain in computational time when using a Lévy mutation scheme instead of a purely local mutation.

**Fig. 4.**
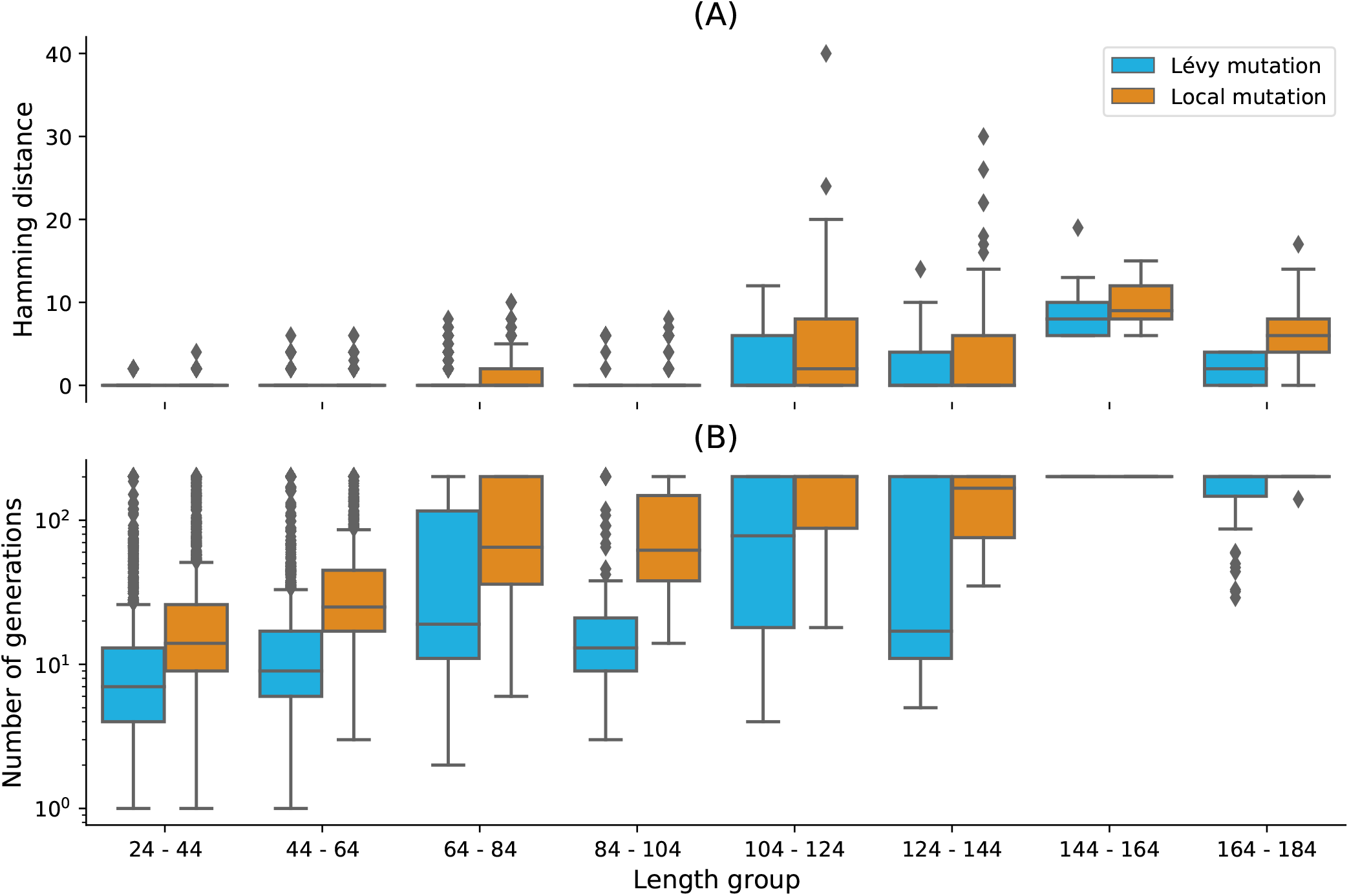
Lévy mutation mode *vs* local mutation (one-point mutation). (A) Hamming distance distributions *vs*. target structure lengths. (B) Number of generations distributions for different length groups. In both (A) and (B), lower values indicate better performance. The target structures are solvable in less than 100 generations for both mutation schemes and most length groups. Still, the difference in the number of generations gets more significant as the target lengths increase, except for the two last length groups for which both mutation schemes mostly failed. The highest difference in terms of median number of generations is 150 for target lengths in the range [124 − 144] (respectively 123, 49, 46, 16, 7, 0, 0 for the length ranges [84 − 104], [64 − 84], [104 − 124], [44 − 64], [24 − 44], [144 − 164], [164 − 184]). Averaging over all length groups, the median number of generations difference between the Levy mutation and the one point mutation is 48 generations.

### Performance on PseudoBase++: aRNAque *vs*. antaRNA

We also compared the sequences designed using `aRNAque` (with the Lévy mutation scheme) to those produced by `antaRNA`. Figures 5A and 5C show the base pair distance distribution for each category of pseudoknotted target structure and the mean of the base pair distance plotted against the length of the target secondary structures. For `antaRNA`, and when using `IPknot` as a folding tool, finding sequences that fold into the target becomes increasingly difficult with pseudoknot complexity (median base-pair distance distribution increases). On the other hand, `aRNAque`’s performance improves as pseudoknot complexity increases (e.g. the mean base-distance decreases with the pseudoknot complexity). In sum, as target length increases, the performance of `antaRNA` (local search) is considerably degraded, while `aRNAque` (Lévy flight search) stays almost constant. A second benchmark using `HotKnots` as a folding tool was performed on the same dataset. For both `aRNAque` and `antaRNA`, the more complex the pseudoknot motifs, the worse is the tool performance (median of the basepair distance distribution increases). Figures 5B and 5D show the base pair distance distributions with respect to the pseudoknot motifs for both `aRNAque` and `antaRNA`. Even though both performances degrade as target length increases, `aRNAque` (Lévy flight evolutionary search) performance remains almost constant for all the target lengths greater than 60.

**Fig. 5.**
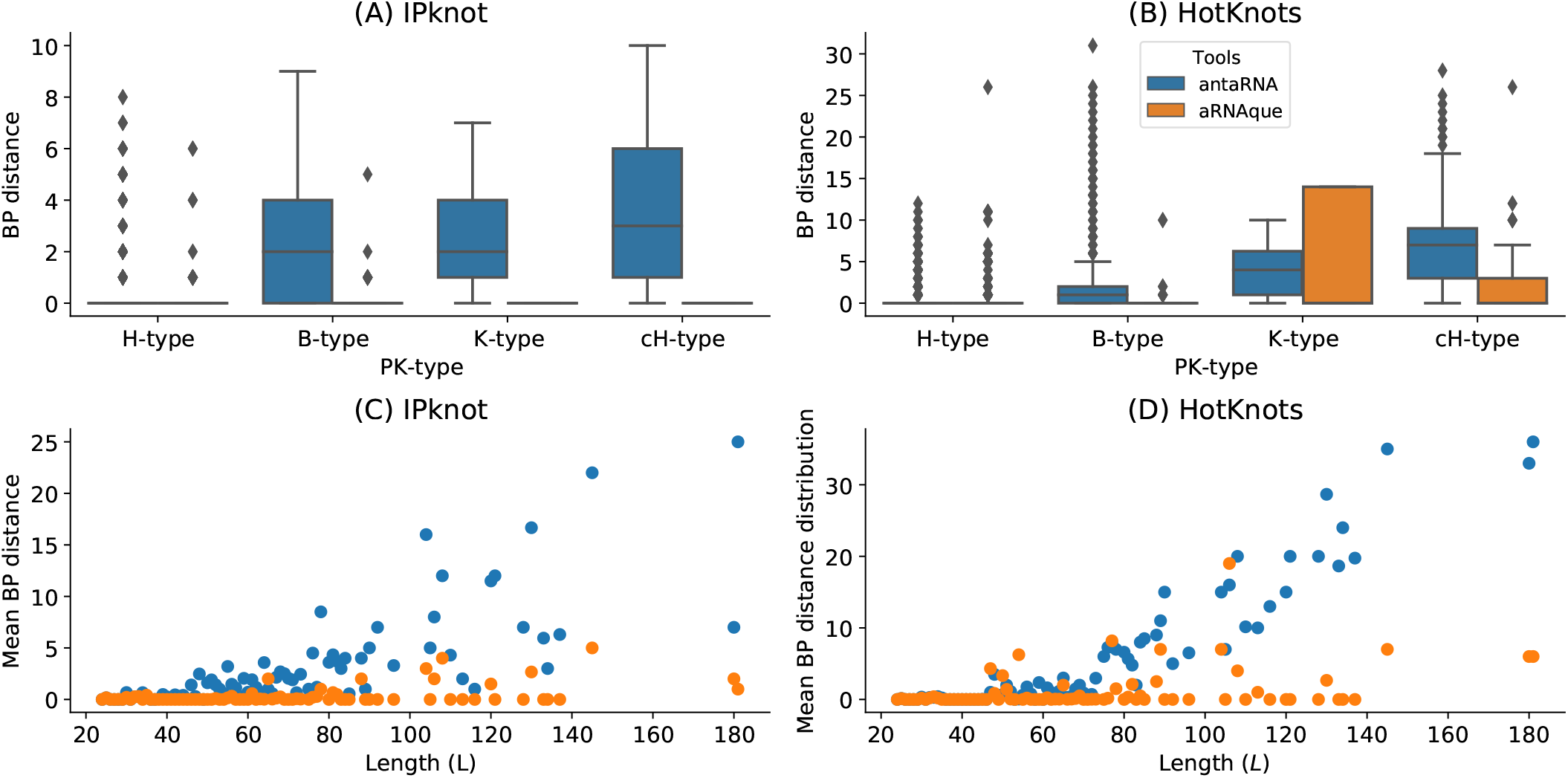
`aRNAque` *vs* `antaRNA` on `PseudoBase++` dataset using both `IPknot` and `HotKnots`. Lower values imply better performance. (A, B) Base pair distance distributions of the designed sequences to the target structure for different pseudoknot types. (C,D) Mean base pair distance against target lengths.

### Performance on Eterna100 dataset

Finally, we performed a third benchmark on the `Eterna100` datasets. First, on the `Eterna100-V1` dataset, the Lévy flight version of `aRNAque` successfully designed 89% of the targets and the one-point mutation (local mutation) version achieved 91% of success, suggesting that for some target structures, local mutation can outperform the Lévy mutation scheme. Combining the two datasets, `aRNAque` solved in total 92% of the targets of `Eterna100-V1` (see also (23)).

When analysing the performance of Lévy flight for low and high base pair densities separately, the median number of generations of high base pair density targets was lower than the one with low base-pair density (8 generations for high density and 18 for the low base pairs density targets). The same observation was drawn for the success rate. For the low base-pair density targets, the Lévy flight achieved 87% (49*/*56) success whereas, for the high base-pair density, it achieved 91% (40*/*44). The same analysis can be done when comparing the one-point mutation results for the high-density targets to the Lévy flight mutation. The median number of generations for the low-density targets when using a onepoint mutation operator was 34 (respectively 24 for the high base pair density targets) (see Figure 7A).

A new benchmark was performed on `Eterna100-V2` with `aRNAque` achieving a 93% success rate. Compared to recently reported benchmark results (34), `aRNAque` achieved similar performance to `NEMO` on `Eterna-V2`: one target was unsolved by all existing tools and one target solved only by `NEMO` remained unsolved by `aRNAque`.

## Discussion

In this work, we provide an updated version of `aRNAque` implementing a Lévy flight mutation scheme that supports pseudoknottted RNA secondary structures. A Lévy mutation scheme offers a compromise between exploration at different scales (mostly local search combined with rare big jumps). Such a scheme significantly improves the number of evaluations needed to hit the target structure, while better avoiding getting trapped in local optima. The benefit of a Lévy flight over a purely local (binomial with *μ <<* 1 or a single point mutation) mutation search allowed us to explore RNA sequence space at all scales. Such a heavy tailed distribution in the number of point mutations permitted the design of more diversified sequences and reduced the number of evaluations of the evolutionary algorithm implemented in `aRNAque`. The main advantage of using a Lévy flight over local search is a reduction in the number of generations required to reach a target. This is because the infrequent occurrence of a high number of mutations allow a diverse set of sequences among early generations, without the loss of robust local search. One consequence is a rapid increase in the population mean fitness over time and a rapid convergence to the target of the maximally fit sequence. To illustrate that advantage, we ran `aRNAque` starting from an initial population of unfolded sequences, both for a “one point mutation” and “Lévy mutation”.

Figures 6A and 6B show respectively the max/mean fitness over time and the number of distinct structures discovered over time plotted against the number of distinct sequences. When using a Lévy mutation scheme, the mean fitness increases faster in the beginning but stays lower than that using local mutations. Later in the optimisation, a big jump or high mutation on the RNA sequences produces structures with fewer similarities and, by consequence, worse fitness. In the (5 − 10)^*th*^ generation, sequences folding into the target are already present in the Lévy flight population, but only at the 30^*th*^ generation are similar sequences present in the local search population. The Lévy flight also allows exploration of both the structure and sequence spaces, providing a higher diversity of structures for any given set of sequences (Figure 6B). Using the mean entropy of structures as an alternate measure of diversity, we see in Figures 6C and 6D how a Lévy flight achieves high diversity early in implementation, and maintains a higher diversity over all generations than a local search algorithm. Although the mutation parameters *P*_*C*_ and *P*_*N*_ influence the absolute diversity of the designed sequences, the Lévy flight always tends to achieve a higher relative diversity than local search, all else being equal.

**Fig. 6.**
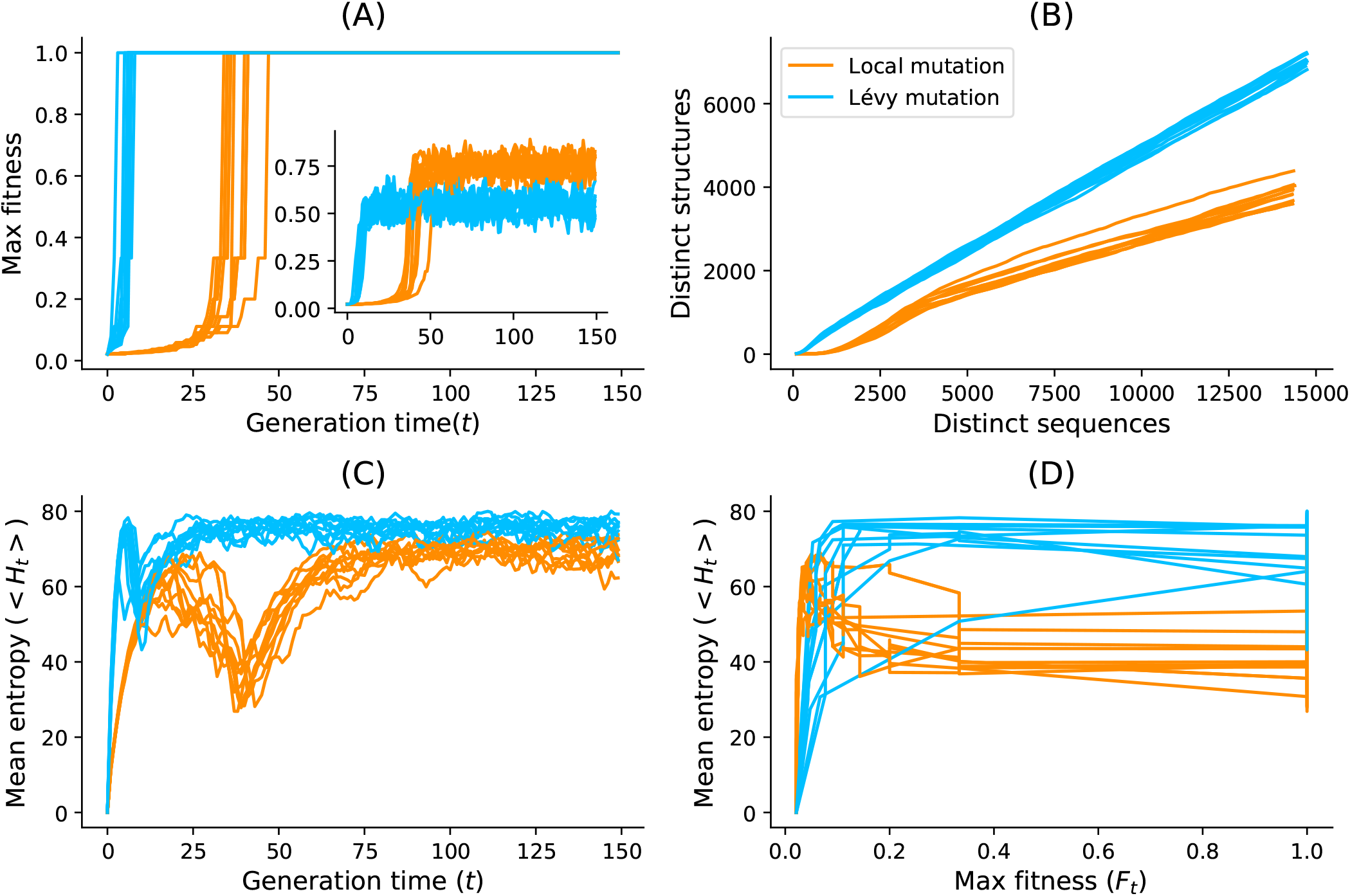
Lévy mutation *vs* one-point mutation: diversity analysis. For the `Eterna100` target structure *[CloudBeta] 5 Adjacent Stack Multi-Branch Loop*, ten independent runs were performed in which a minimum of 10 sequences were designed per run. (A) Max fitness and mean fitness (inset) over time. (B) Distinct sequences *vs*. Distinct structures over time. (C) Mean Shannon entropy of the population sequences over time for both binomial and Lévy mutation. (D) The max fitness plotted against the entropy over time.

We argue that the improved performance of the Lévy Flight over local search in target RNA structures is due to the high base pair density of pseudoknotted structures. Given that pseudoknots present a high density of interactions, there are dramatic increases in possible incorrect folds and thus increasing risk of becoming trapped near local optima (36). Large numbers of mutations in paired positions, as implied by a heavy tailed distribution, are necessary to explore radically different solutions.

To illustrate that Lévy Flight performance was due to base pair density, we clustered the benchmark datasets into two classes: one cluster for target structures with low base pair density (density ≤0.5) and a second cluster for structures with high base pair density (density > 0.5). Figure 7B shows the number of target sequences available in each low and high density category. The number of targets available in each category are colored according to the percentage of pseudoknot-free targets (`Eterna100-V1`) vs. targets with pseudoknots (`Pseudobase++`), showing that pseudoknots are strongly associated with high base pair densities: 71% of the pseudoknotted target structures have a high base pair density. In contrast, the `Eterna100` dataset without psuedoknots has somewhat higher representation at low base pair density. If it is true that improved Lévy Flight performance is indeed tied to base pair density, it is possible that similar heavy-tailed mutation schemes could offer a scalable solution to even more complex inverse folding problems.

**Fig. 7.**
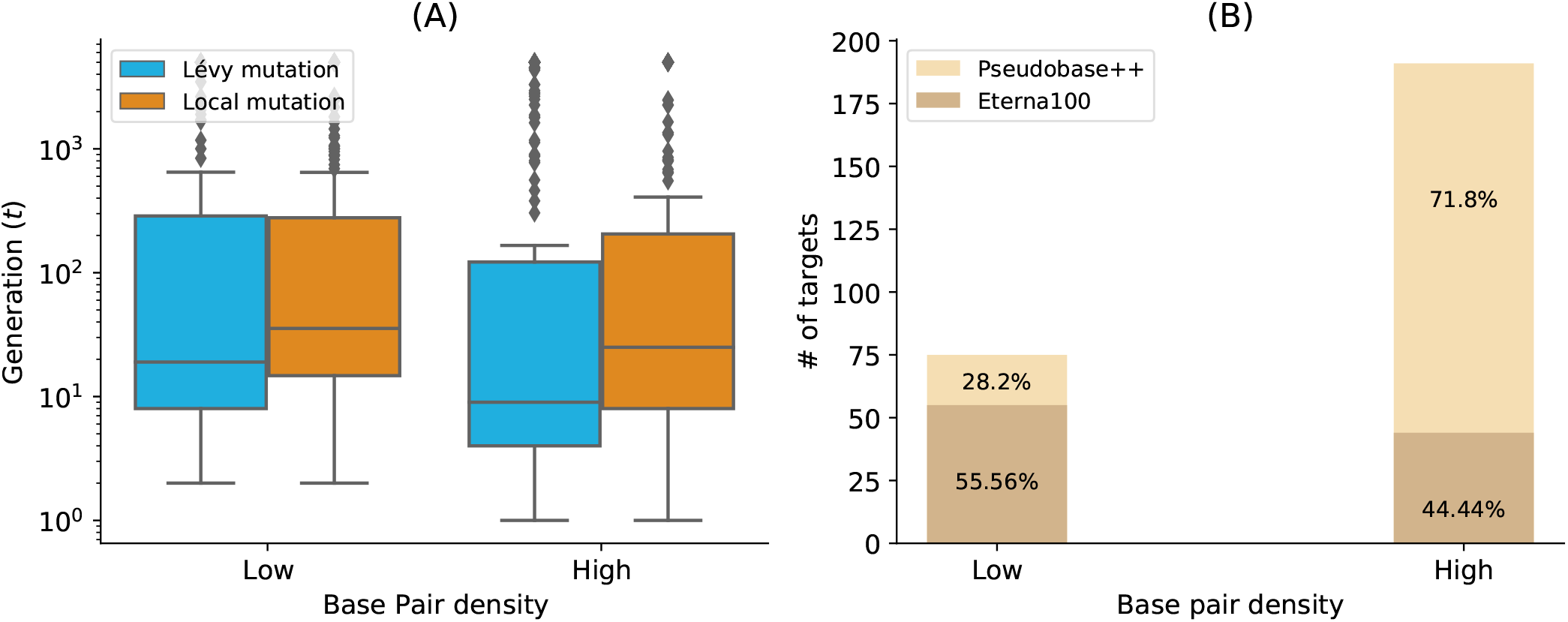
Lévy mutation *vs*. Local mutation: performance analysis with respect to the base-pair density. The higher the base-pair density is, the more useful the Lévy mutation scheme to speed up the optimization EA. (A) Distributions of number of generations for the low and high base-pair density targets of the `Eterna100` dataset. (B) Percentages of targets with low and high base-pair density for the `Eterna100` and `PseudBase++`

Although we believe that Lévy flight-type search algorithms offer a valuable alternative to local search, we emphasise that its enhanced performance over say `antaRNA` is partially influenced by the specific capabilities of existing folding tools. Their limitations may account for the degradation of these tools as the pseudoknot motifs get increasingly complex. Another possible limitation is the fact that most target structures were relatively easy to solve (in less than 100 generations), which possibly allowed local search to perform better than Lévy search in some cases. Further research on more challenging target structures will improve our understanding of which conditions favour local *vs*. Lévy search.

## Conclusion

Our results show general and significant improvements in the design of RNA secondary structures compared to the standard evolutionary algorithm mutation scheme with a mutation parameter ≈1*/L*, where *L* is the sequence solution length. Not only does Lévy flight mutations lead to greater diversity of RNA sequence solutions, but it also reduces the evolutionary algorithm’s number of evaluations, thus improving computing time.

## Availability

The implementation in `python3.7` of `aRNAque` and the benchmark data used in this manuscript are available at https://github.com/strevol-mpi-mis/aRNAque. We also provide the scripts used for the figures and the designed sequences analysis.

## Author’s contributions

Nono S. C. Merleau contributed in the investigation, software coding, visualization, writing – original draft, and writing – review & editing. Matteo Smerlak contributed in the funding acquisition, investigation, methodology, Supervision, validation and the writing – review & editing

## Consent for publication

All authors agree with its contents and with this submission.

## Acknowledgements

We thank the Structure of Evolution Group at the Max Planck Institute for Mathematics in the Sciences, and especially Vincent Messow, Opuu Vaitea and Ian Hatton for useful discussions and insightful comments. The Alexander von Humboldt Foundation provided funding for this work in the framework of the Sofja Kovalevskaja Award endowed by the German Federal Ministry of Education.

